# Ctdnep1 and Eps8L2 regulate dorsal actin cables for nuclear positioning during cell migration

**DOI:** 10.1101/2020.02.20.957761

**Authors:** Francisco J. Calero-Cuenca, Daniel S. Osorio, Sreerama Chaitanya Sridhara, Yue Jiao, Jheimmy Diaz, Sofia Carvalho-Marques, Bruno Cadot, Edgar R. Gomes

## Abstract

Cells actively position their nuclei within the cytoplasm for multiple cellular and physiological functions. Different cell types position their nuclei away from the leading edge to migrate properly. In migrating fibroblasts, nuclear positioning is driven by dorsal actin cables connected to the nuclear envelope by the LINC complex on Transmembrane Actin-associated Nuclear (TAN) lines. How dorsal actin cables are organized to form TAN lines is unknown. Here, we report a role for Ctdnep1/Dullard, a nuclear envelope phosphatase, and the actin regulator Eps8L2, on nuclear positioning. We demonstrate that Ctdnep1 and Eps8L2 directly interact to regulate the formation and thickness of dorsal actin cables required for TAN lines engagement for nuclear positioning. Our work establishes a novel mechanism to locally regulate actin at the nuclear envelope for nuclear positioning.

## Introduction

Positioning the nucleus within the cell is a crucial feature for several cellular events, such as cell migration or asymmetric cell division. Numerous cell types such as epithelial cells, immune cells and multinucleated myofibers precisely position their nuclei for specific functions [1–3]. Consequently, nuclear mispositioning is usually associated to cell dysfunction and disease, from muscular disorders to cancer metastasis [4–7]. The nucleus is permanently under tension from different intracellular forces that result from nucleo-cytoskeletal connections and regulate nuclear movement and mechanotransduction [8–10]. These connections are mainly mediated by the Linker of Nucleoskeleton and Cytoskeleton (LINC) complex, located at the nuclear envelope [11,12]. The LINC complex is composed of KASH domain proteins (Nesprins in mammals) at the outer nuclear membrane that bind to SUN domain proteins, located at the inner nuclear membrane. KASH domain proteins interact with actin, microtubules and intermediate filaments whereas SUN domain proteins interact with the nuclear lamina thus leading to the connection of the nucleus to the cytoskeleton [13,14]. In the last years, the LINC complex was shown to be essential for nuclear dynamics and mechanotransduction [15–18].

The position and the mechanics of the nucleus is especially important during cell migration [19,20]. Different cell types such as fibroblasts or most cancer cells position their nuclei at the cell rear during migration creating a leading edge-centrosome-nucleus axis [21–24]. This cell polarization has been extensively studied in migrating fibroblasts where an actin retrograde flow originated at the leading edge drives dorsal actin cables movement away from the leading edge. The dorsal actin cables connect to the LINC complex at the nuclear envelope creating linear arrays of nuclear envelope proteins called TAN lines (Transmembrane Actin-associated Nuclear lines) resulting in rearward nuclear movement and centrosome reorientation towards the leading edge [22,25,26]. Importantly, dorsal actin cables are distinct from the perinuclear actin cap. The perinuclear actin cap comprises actin bundles on the dorsal side of the nucleus that are attached to focal adhesions. Furthermore, perinuclear actin cap is involved in nuclear shape and mechanotransduction but not in nuclear movement [25,27–32].

During the past years, a set of accessory proteins to the LINC complex that regulate TAN lines establishment and dynamics was identified [33–37]. However, how dorsal actin cables are formed and regulated to form TAN lines is unknown.

In this work, we identified Ctdnep1 and Eps8L2 as main regulators of dorsal actin cables organization and nuclear positioning in migrating cells. Ctdnep1 (C-Terminal Domain Nuclear Envelope Phosphatase 1, also known as Dullard) is a transmembrane Ser/Thr phosphatase found at the nuclear envelope implicated in neuronal development, lipid metabolism and nuclear membrane biogenesis [38–43]. Interestingly, mutations in *CTDNEP1* that result in the loss of wild type allele have been associated to medulloblastoma progression, the most common type of primary brain tumour in childhood [44,45]. Eps8L2 (Epidermal Growth Factor Receptor Kinase Substrate 8-Like Protein 2) is a member of the Eps8-related proteins family. This family is characterized by a C-terminal actin-binding domain with actin capping and bundling activity [46–48]. Their function has been revealed important for filopodia and stereocilia formation as well as cell migration by Rac1 activation [48–51]. Here we described a novel mechanism to locally regulate actin organization for nuclear movement by a nuclear envelope protein. We show that Ctdnep1 and Eps8L2 directly interact to control dorsal actin filaments formation and thickness in the perinuclear region required for TAN lines formation and nuclear positioning in migrating fibroblasts.

## Results

### Ctdnep1 and Eps8L2 are required for nuclear positioning and centrosome reorientation in migrating fibroblasts

The LINC complex is the main player connecting the nucleus to the cytoskeleton and it has an essential role for nuclear movement and positioning [10,11,33]. In order to identify new regulators of nuclear position we tested the involvement of Ctdnep1, a nuclear envelope and endoplasmic reticulum Ser/Thr phosphatase in polarization of migrating cells [40,52].

We used small interference RNA (siRNA) to deplete Ctdnep1 in 3T3 fibroblasts grown to confluence and serum-starved for 48 hours (Figure S1A). We wounded the monolayer and stimulated cell polarization by adding LPA (Figure 1A) as previously described [22,53]. We quantified nuclear and centrosome positions relative to the cell centroid as well as percentage of centrosome reoriented cells as a readout of cells polarization (Figure 1B-C). Cells treated with Control siRNA positioned their nuclei away from the cell centroid upon LPA stimulation whereas the nucleus of non-LPA treated cells remained near the cell centroid. When the LINC complex was disrupted by Nesprin2G depletion, nuclear positioning away from the cell centroid was inhibited (Figure 1B) and centrosome did not reorient (Figure 1C), as previously showed [25,33]. Interestingly, upon depletion of Ctdnep1 using two different siRNA oligos, nuclear positioning away from the cell centroid and centrosome reorientation were also inhibited (Figure 1A-C), thus suggesting a role for Ctdnep1 on nuclear positioning.

**Figure 1.**
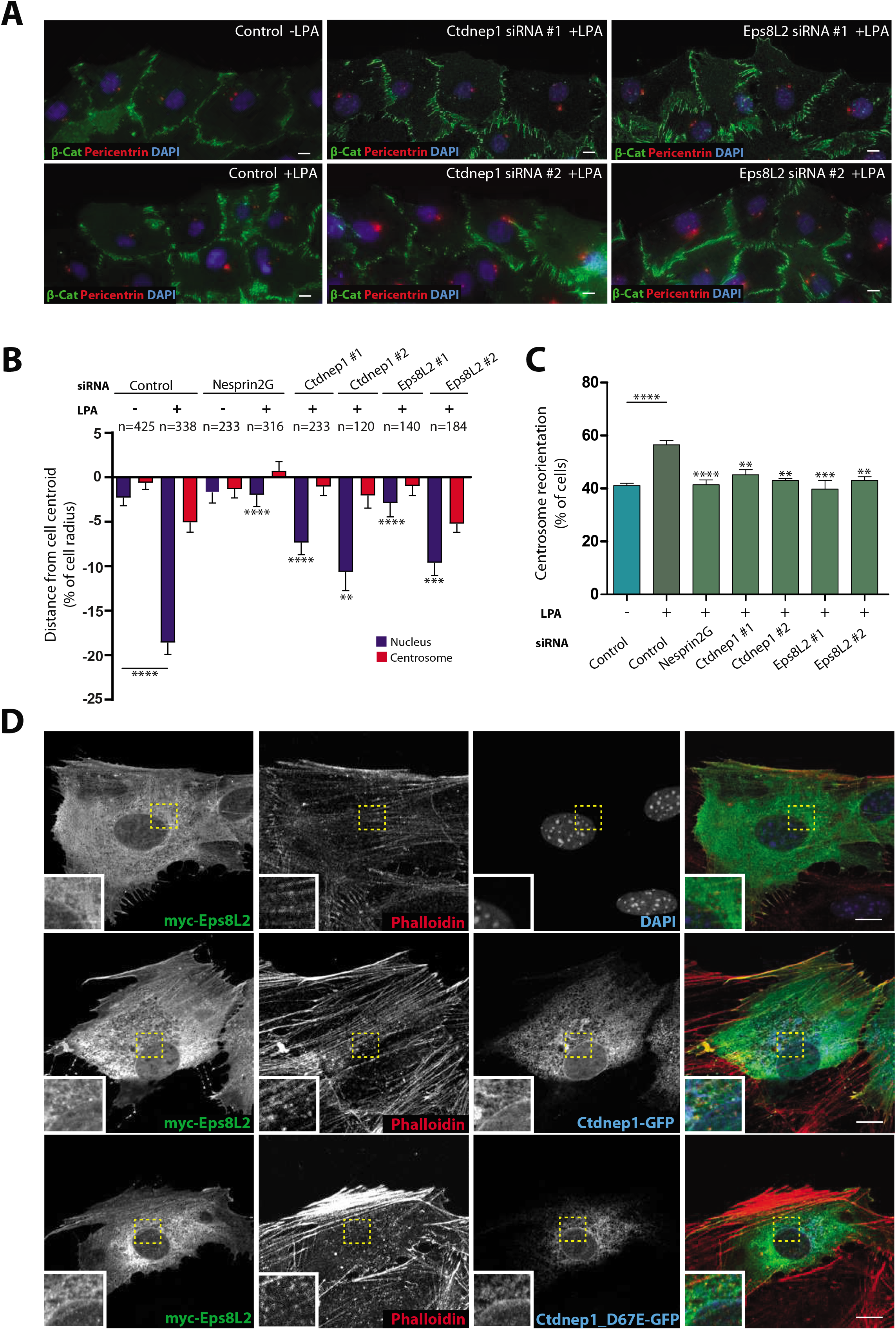
Ctdnep1 and Eps8L2 are required for nuclear positioning in migrating fibroblasts. (**A**) Representative images of wound-edge 3T3 fibroblasts with or without LPA stimulation in Control, Ctdnep 1 and Eps8L2 siRNAs stained for β-catenin (green, cell contacts), pericentrin (red, centrosome) and DAPI (blue, nucleus). (**B**) Average positions of the nucleus (blue) and centrosome (red) relative to the cell centroid in cells treated with Control, Nesprin2G, Ctdnep 1 and Eps8L2 siRNAs. (**C**) Percentage of oriented cells in the conditions analyzed in B. (**D**) Representative images of wound-edge wildtype fibroblasts stimulated with LPA and microinjected with myc-Eps8L2, Ctdnep 1-GFP and Ctdnep 1_D67E-GFP stained for myc (green, Eps8L2), phalloidin (red, Actin) DAPI (blue, nucleus) or GFP (blue, Ctdnep 1 and Ctdnep l_D67E). The bottom left inset is a zoom in of the area indicated by the yellow square. Scale bars: 10 μm. Data are represented as mean +/− SEM.

To identify the molecular mechanism by which Ctdnep1 regulates nuclear positioning, we performed an Yeast Two-Hybrid (Y2H) screen using the cytoplasmic domain of Ctdnep1 as bait to identify potential Ctdnep1 interacting partners. Among different hits, we identified Eps8L2, an actin-binding protein member of Eps8 family (Figure S1C). Eps8 proteins are responsible for actin remodelling mediated by RTK-activated signalling pathway [48]. In order to address the role of Eps8L2 in nuclear positioning, we depleted Eps8L2 from fibroblasts using two different siRNA oligos (Figure S1B). Transient depletion of Esp8L2 reduced nuclear positioning away from the cell centroid (Figure 1A-B) as well as centrosome reorientation (Figure 1C) similarly to Ctdnep1 depleted fibroblasts. These results indicate that Ctdnep1 and Eps8L2 are required for nuclear positioning as well as cell polarization in migrating fibroblasts.

### Ctdnep1 interacts directly with Eps8L2 independently of Ctdnep1 phosphatase activity

Taking into account that Ctdnep1 and Eps8L2 showed a positive interaction in the Y2H assay and are both required for nuclear positioning, we analyzed the subcellular localization of Ctdnep1 and Eps8L2 in migrating fibroblasts. Due to the lack of suitable antibodies for immunohistochemistry, we performed microinjection of cDNA encoding Ctdnep1. Ctdnep1_D67E (a mutant with no phosphatase activity [42]) and Eps8L2 in wound edge cells to allow a tight control of protein expression. Ctdnep1 localiz es to the nuclear envelope and endoplasmic reticulum independently of its phosphatase activity (Figure 1D and Figure S2A). Using digitonin permeabilization we also observed Ctdnep1 is present at the outer nuclear membrane (Figure S2B). On the other hand, Eps8L2 localizes in the cytoplasm and it is enriched at the leading edge, actin filaments, cell projections and perinuclear region (Figure 1D). Furthermore, we observed partial colocalization of both Ctdnep1 and Eps8L2 at the perinuclear area (Figure 1D).

We then investigated if Ctdnep1 physically interacts with Eps8L2 and if this interaction was direct. We performed an *in vitro* pull down using recombinant His-Ctdnep1_Cter, a cytoplasmic version of Ctdnep1 without the transmembrane domain (in order to make the protein soluble, Figure 2A), We found His-Ctdnep1_Cter pulled down GST-Eps8L2 strongly suggesting that both proteins interact directly (Figure 2B). Additionally, we investigated if the Ctdnep1 phosphatase activity was involved in this interaction. We immunoprecipitated endogenous Eps8L2 or Flag-Eps8L2 co-expressed with Ctdnep1-GFP or Ctdnep1_D67E-GFP (the phosphatase-dead mutant) in U2OS cells. We were able to co-immunoprecipitate endogenous and expressed Eps8L2 with both Ctdnep1 constructs (Figure 2C-D) suggesting that the phosphatase activity of Ctdnep1 is not required for the interaction. Overall, these results indicate that the Ctdnep1-Eps8L2 physical interaction is direct and Ctdnep1 phosphatase activity is not required for its interaction with Eps8L2.

**Figure 2.**
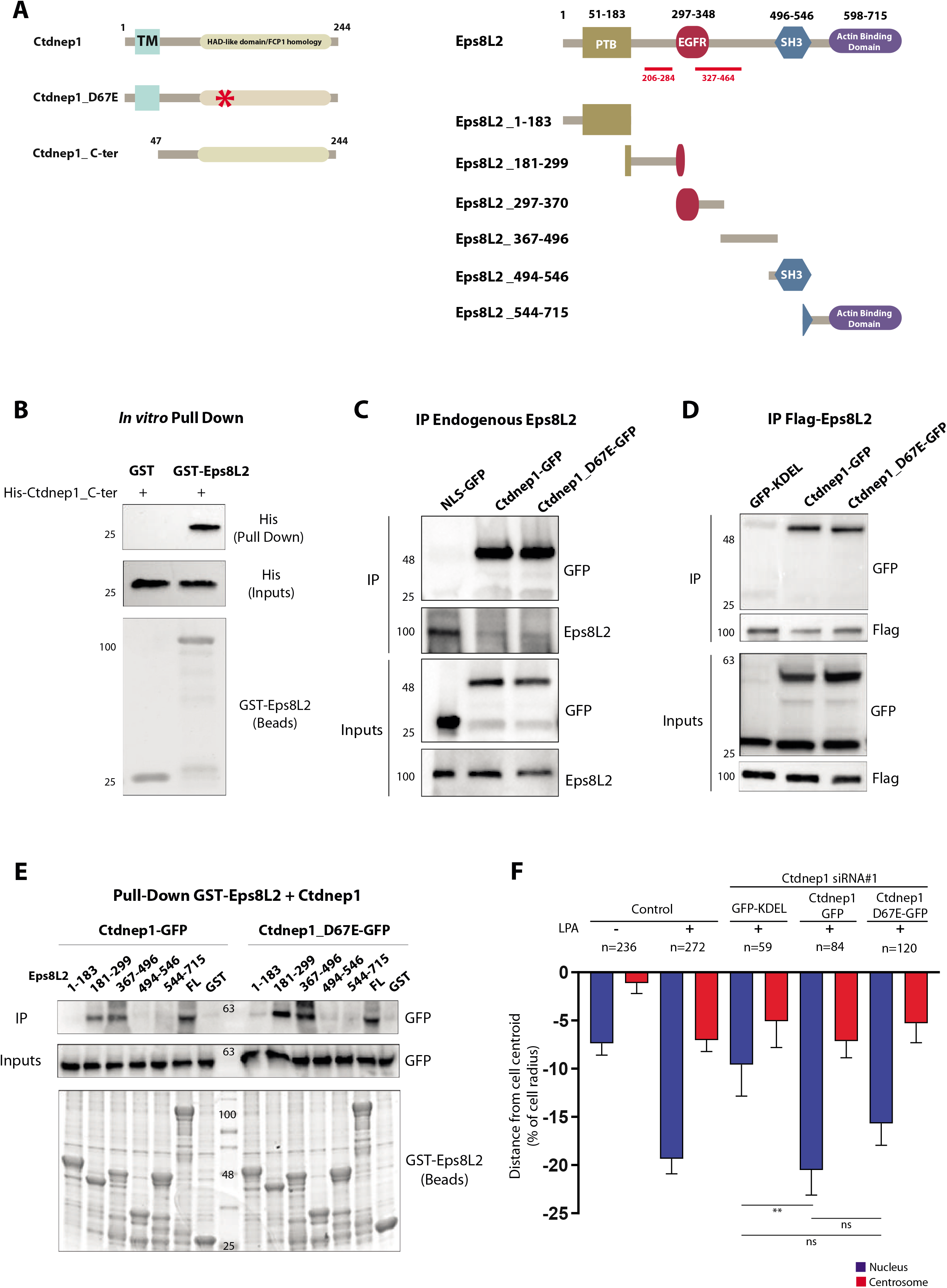
Ctdnep1 and Eps8L2 interact directly. (**A**) Schematic representation of Ctdnep1, Ctdnep1_D67E (no phosphatase activity, red asterisk denotes D67E mutation), Ctdnep1_C-ter and Eps8L2 proteins showing their different protein domains and fragments used. (**B**) *in-vitro* pull-down of recombinant GST-Eps8L2 bound to glutathione agarose beads with purified recombinant His-Ctdnep1_C-ter (Ctdnep1 without the transmembrane domain). (**C**) Co-Immunoprecipitation of endogenous Eps8L2 from SRKB cells with GFP-NLS, Ctdnep1-GFP and Ctdnpel_D67E-GFP overexpressed in U2OS cells. (**D**) Co-Immunoprecipitation of Flag-Eps 8L2 overexpressed in U2OS cells and posteriorly incubated with GFP-KDEL, Ctdnep1-GFP and Ctdnep1_D67E-GFP overexpressed in U2OS cells (**E**) Pull-down assay of recombinant GST-Eps8L2 and its different fragments bound to glutathione agarose beads with Ctdnep1-GFP and Ctdnep1_D67E-GFP overexpressed in U2OS cells. (**F**) Average positions of the nucleus (blue) and centrosome (red) in cells treated with Ctdnep1 siRNA and microinjected with KDEL-GFP, Ctdnep1-GFP and Ctdnep1_D67E-GFP. Data are represented as mean +/− SEM.

### Ctdnep1 and Eps8L2 regulate nuclear positioning independently of dephosphorylation

Eps8L2 contains four described domains: PTB, EGFR, SH3 and an Actin-Binding Domain at the C-terminal (Figure 2A) [48,54]. We cloned different fragments of Eps8L2 taking into account these described domains and produced recombinant protein of each fragment tagged with GST (Figure 2A). We then performed pull down assays using extracts of U2OS cells expressing Ctdnep1-GFP or Ctdnep1 _D67E-GFP. We observed that Ctdnep1-GFP and Ctdnep1 _D67E-GFP were pulled down by Eps8L2 fragments 181-299 and 367-496, to a similar extent as full length Eps8L2 (Figure 2E). These regions are located between the PTB and EGFR domains and between the EGFR and SH3 domains respectively and overlap with the two regions detected in our Yeast Two-Hybrid screen (Figure 2A).

Since both Ctdnep1 and Ctdnep1 phosphatase dead mutant (D67E) were able to bind to Eps8L2, we wondered if the phosphatase activity of Ctdnep1 was required for nuclear positioning. We microinjected siRNA resistant Ctdnep1-GFP and Ctdnep1 _D67E-GFP as well as GFP-KDEL as a control, in starved wound edge fibroblasts transfected with Ctdnep1 siRNA. We observed that upon LPA stimulation, Ctdnep1-GFP fully restored nuclear positioning away from the cell centroid whereas Ctdnep1 _D67E-GFP partially rescued this nuclear positioning (Figure 2F). Therefore, the phosphatase activity of Ctdnep1 does not seem to be involved on nuclear movement.

It was previously reported that Eps8 function during axonal filopodia formation is regulated by phosphorylation [50] and specifically for Eps8L2, different reports identified several amino acids that can be phosphorylated [55–59]. Therefore, we tested if Eps8L2 was a Ctdnep1 substrate. We performed Phos-tag SDS-page to analyzed Eps8L2 phosphorylation in SKBR3 cell lysates (which have high levels of endogenous Eps8L2 [60]) treated with Control and Ctdnep1 siRNAs. When we treated the lysates with Lambda phosphatase (to dephosphorylate all proteins in the sample), Eps8L2 band appeared in a lower molecular weight than without Lambda treatment (Figure S3 A). This result suggests Eps8L2 is phosphorylated as previously described. However, we did not observe any differences between Control or Ctdnep1 siRNAs (Figure S3 A). Additionally, we performed mass spectrometry to identify phosphorylation sites in myc-Eps8L2 purified from U2OS cells cotransfected with Ctdnep1-GFP or Ctdnep1_D67E-GFP. We identified several phosphorylated sites in Eps8L2, some of them were previously reported. However, we did not observe any difference regarding the phosphorylation profile related to the phosphatase activity of Ctdnep1 when we compared the different conditions (Figure S3B). Therefore, our results support a dephosphorylation-independent role of Ctdnep1 and Eps8L2 on nuclear positioning.

### Ctdnep1 and Eps8L2 are required for TAN lines formation

Nuclear movement during centrosome reorientation is driven by actin retrograde flow and requires the formation of TAN lines by the connection of dorsal actin cables to the LINC complex [22,25]. We first measured actin retrograde flow in fibroblasts stably expressing Lifeact-mCherry upon depletion of Ctdnep1 and Eps8L2. We observed that the percentage of cells with actin retrograde flow was slightly decreased in cells depleted for Ctdnep1 and Eps8L2 when compared to Control (Figure S4A). Interestingly, the speed of the actin retrograde flow near the leading edge and on top of the nucleus, where the TAN lines are formed, was slightly increase in Eps8L2 depletion whereas was not altered in Ctdnep1 depletion (Figure S4B). Therefore, the minor changes we observed on actin retrograde flow upon Ctdnep1 and Eps8L2 depletion cannot account for the observed effect on nuclear movement.

We then explored the role of Ctdnep1 and Eps8L2 in TAN lines dynamics. First, we evaluated if Ctdnep1 or Eps8L2 were enriched at TAN lines. To address this question we microinjected GFP-miniNesprin2G (miniN2G), a functional reporter of TAN lines [25], together with Ctdnep1-Flag. Ctdnep1_D67E-Flag and myc-Eps8L2 in wild type wound edge fibroblasts. Upon LPA stimulation we observed that Eps8L2, but not Ctdnep1 or Ctdnep1_D67E, co-localized with dorsal actin cables at TAN lines (Figure 3A). We then tested whether Ctdnep1 or Eps8L2 were required for TAN lines formation. We microinjected GFP-miniN2G in wound edge fibroblasts transfected with Control, Ctdnep1 or Eps8L2 siRNAs, followed by LPA stimulation. Depletion of both Ctdnep1 or Eps8L2 reduced the number of cells with TAN lines (Figure 3B-C). In addition, the average number of TAN lines per cell was dramatically decreased in cells depleted for Ctdnep1 or Eps8L2 (Figure 3B-D). Our results suggest that Ctdnep1 and Eps8L2 are involved in TAN lines formation during nuclear positioning in migrating fibroblasts.

**Figure 3.**
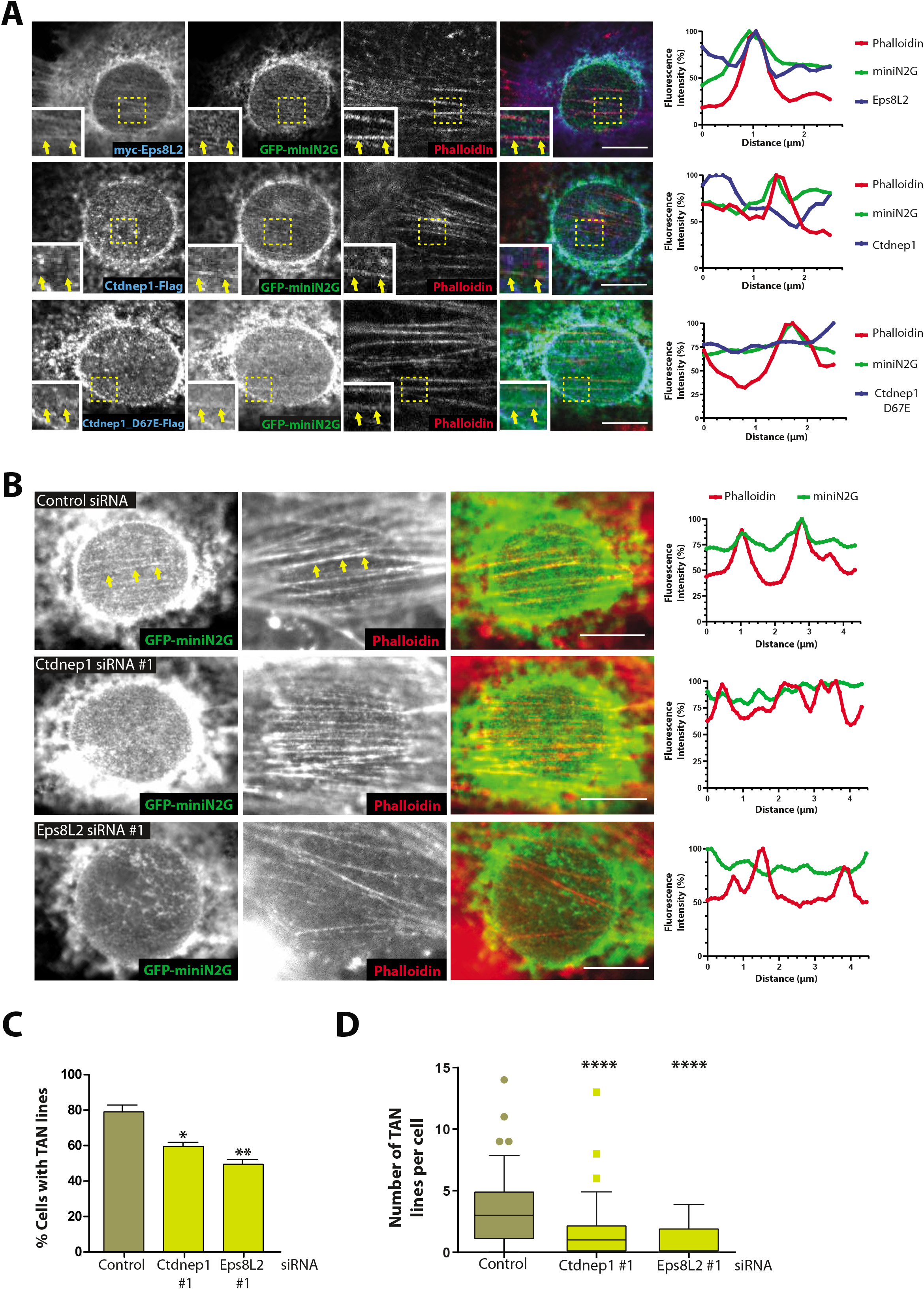
Ctdnep1 and Eps8L2 affect TAN lines formation. (**A**) Wound-edge fibroblast stimulated with LPA and microinjected with GFP-miniN2G and myc-Eps8L2 (top panel) Ctdnep1-Flag (middle panels) and Ctdnep1_D67E-Flag (bottom panel). Cells were stained for GFP (green), phalloidin (red), myc (blue) and flag (blue). Colocalization of miniN2G and Actin (Phalloidin) in linear arrays at the nuclear envelope is indicated by yellow arrows in the insets. Eps8L2, but not Ctdnep1, colocalize with actin at TAN lines. Line scans for each channel of the TAN lines marked by the yellow arrows are represented in the right plots. (**B**) Wound-edge fibroblasts transfected with Control, Ctdnep1 and Eps8L2 siRNAs were microinjected with GFP-miniN2G to analyse TAN lines formation. Cell were stained for GFP (green) and phalloidin (red). TAN lines can be visualized in the Control (yellow arrows). Line scans for each channel are represented in the right plots. Scale bar: 10 μm. (**C**) Quantification of the percentage of cells presenting at least one TAN line in the conditions described in B. Data are represented as mean +/− SEM. (**D**) Quantification of number of TAN lines per cell in the conditions described in B. Data are represented as 5-95 percentile values.

### Ctdnep1 and Eps8L2 regulate dorsal actin organization

TAN line formation requires actin retrograde flow and the formation of dorsal actin cables [25]. Furthermore, we showed that Eps8L2 localizes to dorsal actin cables on TAN lines (Figure 3A) and both Ctdnep1 and Eps8L2 are required for TAN lines formation (Figure 3B, C and D). Therefore, we investigated if Ctdnep1 and Eps8L2 were specifically involved in the formation of dorsal actin cables. We examined dorsal actin organization in serum-starved wounded monolayer of fibroblasts treated with Control, Ctdnep1 and Eps8L2 siRNAs after stimulation with LPA. We observed a decrease of the number of dorsal actin cables in cells depleted for Eps8L2 (Figure 4A-B), without any changes on focal adhesions (Figure S4C). We then analyzed in more detail the dorsal actin cables that connect to the TAN lines in the absence of Ctdnep1 and Eps8L2 and found a reduction of dorsal actin cables thickness when compared to control siRNA (Figure 4C). Finally, we tested if the phosphatase domain of Ctdnep1 was involved in regulating dorsal actin cables thickness. We microinjected siRNA resistant Ctdnep1-GFP and Ctdnep1 _D67E-GFP as well as GFP-KDEL as control in starved wound edge fibroblasts transfected with Ctdnep1 siRNA. We observed that upon LPA stimulation, Ctdnep1-GFP and Ctdnep1 _D67E-GFP fully restored dorsal actin cables thickness (Figure 4C and Figure S5A). These results support a role for Ctdnep1 and Eps8L2 on the formation and maintenance of dorsal actin cables required for TAN lines formation and nuclear movement.

**Figure 4.**
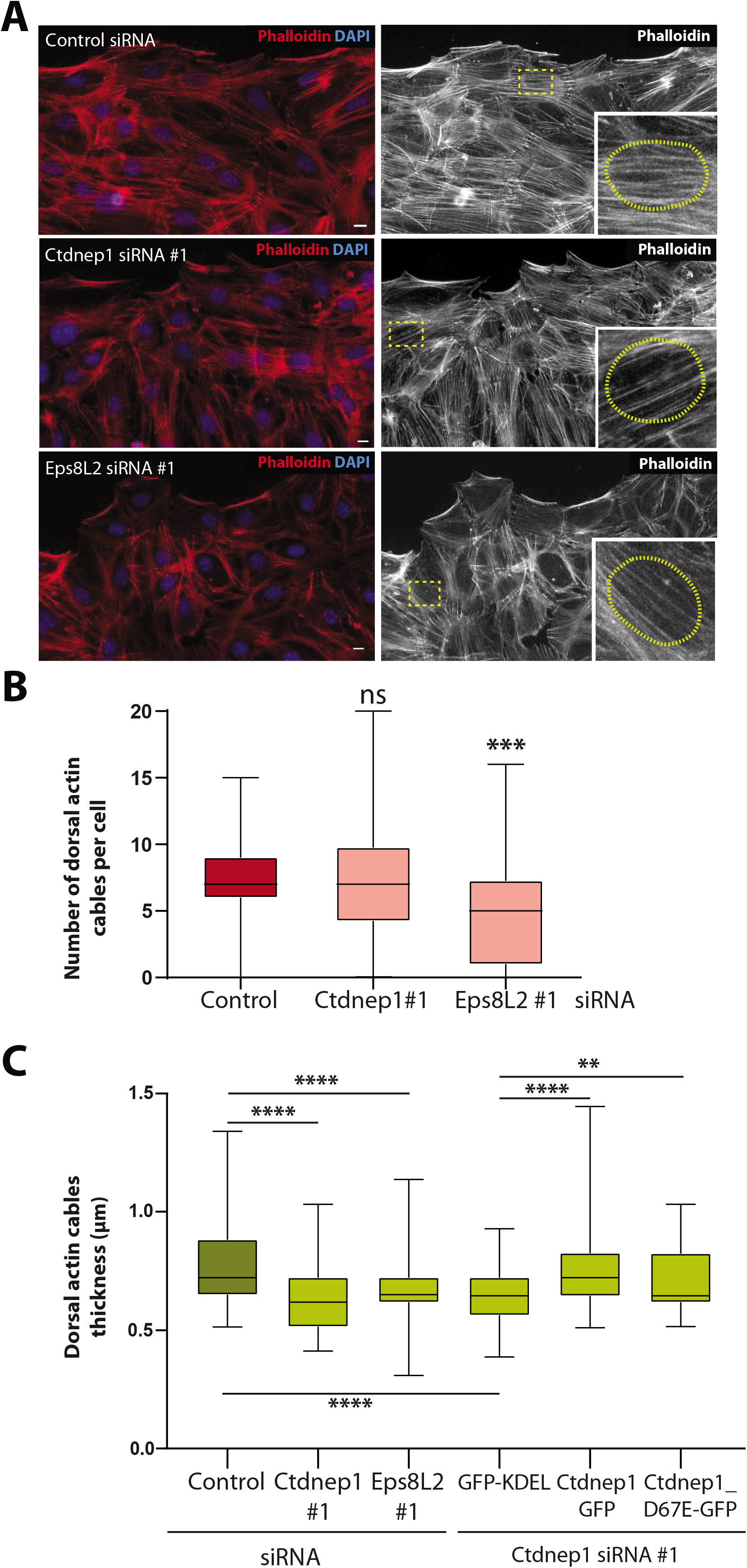
(**A**) Representative images of wound-edge fibroblasts stimulated with LPA and treated with Control, Ctdnep1 and Eps8L2 siRNAs and stained for phalloidin (red, actin) and DAPI (blue, nucleus). The bottom right inset is a zoom in of the area indicated by the yellow square. The yellow dots circles in the insets denote nucleus border. Scale bar: 10 μm. (**B**) Quantification of number of dorsal actin cables on top of the nucleus in the conditions described in A. (**C**) Quantification of dorsal actin cables thickness in the conditions described in A and in fibroblasts transfected with Ctdnep1 siRNA and microinjected with KDEL-GFP, Ctdnep1-GFP and Ctdnep1_D67E-GFP. Data are represented as minimum-maximum values.

## Discussion

Nucleus-cytoskeleton connection has been revealed crucial for nuclear dynamics and cell function. Here we unravel a new mechanism to locally regulate the actin cytoskeleton for nuclear positioning in migrating fibroblasts. We showed that the nuclear envelope protein Ctdnep1 and the actin binding protein Eps8L2 are involved in dorsal actin filaments formation and maintenance required for the engagement of the LINC complex at TAN lines for nuclear movement.

We demonstrated Ctdnep1 and Eps8L2 directly interact. Additionally, we provided evidence for a role of this interaction on nuclear movement, in a dephosphorylation-independent manner. Ctdnep1-Eps8L2 interaction may be transient but essential to regulate Eps8L2 bundling activity stabilizing actin filaments that can properly organize LINC complexes into TAN lines for nuclear movement.

Ctdnep1 depletion does not affect overall actin organization in migrating fibroblast. Conversely, when we reduced Eps8L2 protein levels we observed a dramatic reduction of dorsal actin cables on the top of the nucleus, without affecting focal adhesions and stress fibers, Both Ctdnep1 and Eps8L2 depletions lead to a decrease in dorsal actin cables thickness probably due to lack of Eps8L2 bundling activity. This would impair TAN lines formation due to 1) a faster speed in actin cables retrograde flow and 2) reduced contact surface or traction forces between thinner actin cables and LINC complex. Both scenarios would reduce the ability of actin cables to bind to the LINC complex and to engage TAN lines for nuclear movement.

Ctdnep1 function on lipid metabolism and nuclear membrane biogenesis has been associated with Ctdnep1 phosphatase activity [38,61]. We did not find any role for the phosphatase activity of Ctdnep1 on nuclear movement, interaction with Eps8L2 or dorsal actin cables regulation (Figures 2 and 4). Furthermore, we did not detect any changes in the phosphorylation of Eps8L2 dependent of the phosphatase activity of Ctdnep1 (Figure S3). Therefore, our results suggest a phosphatase independent function for Ctdnep1 on the regulation of dorsal actin cables for nuclear movement.

In the last years, perinuclear actin organization and function have been revealed important for nuclear positioning and cell migration [27,32,62]. Actin organization around the nucleus is crucial for dendritic cells to migrate in 3D environments [62]. Additionally, the LINC complex displays a central role for mechanosensing and mechanotransduction, due to its function as a bridge between the cytoskeleton and the nucleus [15,18,63,64]. Nevertheless, how this LINC complex function is regulated as well as the different cellular processes where it is involved needs to be deciphered. The mechanism of perinuclear actin organization required for nuclear movement described in this report can therefore be involved in the multiple function of LINC complex mediated nucleus-cytoskeleton connection.

## Methods

### Plasmids

Ctdnep1 was directly cloned from an NIH 3T3 mRNA library using the SuperScript^®^ III One-Step RTPCR System (Life Technologies) using primers Ctdnep1 _FL_N1_For and Ctdnep1 _FL_N1_Rev that include AttB recombination site to pDONR201 Gateway donor vectors (Life Technologies). Ctdnep1_C-ter was amplified from Ctdnep1 Full Length (FL) entry vector by using primers Ctdnep1 _Cter_C1_For and Ctdnpe1_FL_C1_Rev. For phosphatase-null point mutant Ctdnep1_D67E, site-directed mutagenesis was performed by PCR amplification of pDONR221 Ctdnep1 FL vector using primers Ctdnep1 _D67E_For and Ctdnep1 _D67E_Rev followed by DpnI endonuclease digestion of the parent (methylated) DNA chain. After sequence confirmation, entry vectors were recombined using the Gateway system with pDEST47 (Life Technologies) for C-terminal GFP-Tag fusions proteins and pDEST17 (a gift from Fanny Jaulin Lab) for N-terminal 6xHis-Tag fusion protein.

Ctdnep1 WT-2xFlag were a gift from Shirim Bahmanyar. Ctdnep1_D67E-2xFlag was generated by site-directed mutagenesis by PCR amplification of Ctdnep1-Flag using the primers hCtdnep1 _D67E_For and hCtdnep1_D67_Rev followed by DpnI endonuclease digestion of the parent (methylated) DNA chain.

Human Eps8L2 cDNA was synthetized (Life Technologies) with attB sites to clone it in pDONR201 Gateway donor vector (Life Technologies). The entry vector generated was recombined using the gateway system with pEZYflag (Addgene #18700), pDEST1 or pRK5mycGW (a gift from Fanny Jaulin Lab) to create Flag, GST or Myc N-terminal fusion proteins. For rescue experiments. Ctdnep1-GFP and Ctdnep1 _D67E-GFP siRNA resistant sequences adding silent mutations for Ctdnep1 siRNAs #1 and #2 were synthesized (GenScript) and cloned directly to pcDNA3.1(+)-C-eGFP vector.

Eps8L2 fragments (1-183, 181-299, 297-370, 367-496, 494-546, 544-715) were amplified from Eps8L2 full length using the primers Eps8L2_S1, Eps8L2_R183, Eps8L2_S181, Eps8L2_R299, Eps8L2_S297, Eps8L2_R370, Eps8L2_S367, Eps8L2_R496, Eps8L2_S494, Eps8L2_R546, Eps8L2_S544, Eps8L2_R715 that include attB sites to clone the different fragments in pDONR201 Gateway donor vector (Life Technologies). The entry vectors generated were recombined using the Gateway system with pDEST15 or pRK5mycGW to GST or Myc N-terminal fusion proteins. GFP-NLS (a gift from Jan Lammerding Lab), GFP-Kdel [65], GFP-miniN2G (a gift from Gregg Gundersen Lab), Lifeact-mcherry (a gift from Olivier Pertz Lab), pGEX-6P-1 (GE Healthcare Life Sciences).

### RT-qPCRs for siRNA validation

After 72h transfection with the different siRNAs, total RNA was extracted from fibroblasts cultures using TRIzol^®^ Reagent (ThermoFisher Scientific) according to the manufacturer’s instructions. RNA yield and purity was assessed using Nanodrop 2000 apparatus, cDNA was synthesized using the High capacity RNA-to-cDNA kit (Applied Biosystems) as manufacturer’s instructions indicate. Reverse transcription quantitative PCR (RT-qPCR) was performed using Power SYBR Green PCR MasterMix (Alfagene) according to the manufacturer’s instructions and using primers forward and reverse at 0,25 μM (final concentration) as well as 1 =20 cDNA dilution. Amplification of Ctdnep1 mRNA was performed using primers Ctdnep1_Ex3_Ms_For and Ctdnep1_Ex3_Ms_Rev. Amplification of Eps8L2 mRNA was performed using primers Eps8L2 Eps8L2_Ex5_Ms_For and Eps8L2_Ex5_Ms_Rev. As a control, housekeeping gene *gapdh* was amplified using the primers Gapdh_Ms_For and Gapdh_Ms_Rev. RNA extraction, cDNA synthesis and RT-qPCRs were performed three times and relative transcription levels were determined using the ΔΔct method.

### Cell culture, siRNA and cDNA infection and microinjection

NIH-3T3 fibroblasts were cultured in DMEM with no sodium pyruvate, 10% bovine calf serum, 10 mM HEPES and penicilin/streptomycin at 500 units/ml. The different siRNAs were transfected as previously described [33] using Lipofectamine RNAiMAX (Invitrogen), Ctdnep1 siRNA #1, Eps8L2 siRNA #1 and Eps8L2 siRNA #2 contain Silencer Select modifications (Life Technologies), Ctdnep1 siRNA #2 and Nesprin2G siRNA were from Genecust Europe, Control siRNA (Silencer Select Negative Control N° 1 siRNA from ThermoFisher Scientific), Microinjections for siRNA rescue and immunofluorescence were performed as described in [22,53] using a Xenoworks microinjection system (Sutter Instruments), A stable cell line expressing Lifeact-mCherry was created to analyse actin retrograde flow by infecting NIH3T3 fibroblasts with lentivirus carrying LifeActin-mCherry produced in HEK 293T cells (pLALI backbone).

U2OS cells were cultured in DMEM with sodium pyruvate, 10% Fetal Bovine Serum, and penicilin/streptomycin at 500 units/ml, Cells were transfected using Lipofectamine 3000 (Invitrogen) with the proper plasmids for 24 hours. After transfection, lysates were made for immunoprecipitation.

SKRB-3 cells were cultured in DMEM with sodium pyruvate, 10% Fetal Bovine Serum, 1% Non-essential amino acids and penicilin/streptomycin at 500 units/ml.

### Centrosome reorientation and nuclear movement analysis

Wound assays were performed as described in [53]. In summary, cells were plate on acid-wash coverslips, at the same time that were transfected with the proper siRNA, in a confluence that allows a cell monolayer in the end of the assay. 36-48 hours after transfection (cell confluence around 50-60%), cells were starved for 48 hours (DMEM with no sodium pyruvate, 10mM HEPES and penicilin/streptomycin at 500 units/ml). Then, the cell monolayer was scratch-wounded with a pipette tip and cells were stimulated with 20mM of LPA for 2 hours. Cells were then fix for immunostaining. For TAN lines analysis, stimulation with 20mM of LPA was performed for 50 minutes. For microinjections, cells were scratch-wounded and microinjected after 48 hours of starvation. Two hours after microinjection, cells were stimulated with 20mM LPA for 2 hours or 50 minutes. Centrosome and nucleus positions were analysed using the software *Cell Plot* developed by Gregg Gundersen Lab (http://www.columbia.edu/~wc2383/software.html).

### Immunofluorescence

Cells were fixed in 4% paraformaldehyde in PBS for 10 minutes at room temperature. Permeabiliz ation with 0.3% Triton X-100 for 5 minutes or 40 μg/ml Digitonin for 3 min at room temperature were posteriorly performed. Primary and secondary antibodies were diluted in PBS containing 10% goat serum. Cells were incubated with primary antibodies for 1 hour at room temperature. After washing three times (20 minutes each) with PBS, cells were incubated with secondary antibodies for 1 hour at room temperature. Cells were washed with PBS (3 times, 20 minutes each time). Fluoromount-G (Invitrogen) was used to prepare the coverslips for cell imaging. The following primary antibodies were used: rabbit anti-β-Catenin 1:200 (Invitrogen #712700), mouse anti-Pericentrin 1:200 (BD-Biosciences # 611814), mouse anti-Flag 1:200 (Sigma-Aldrich #F1804), chicken anti-GFP 1:1000 (Aves Labs #GFP-1020), rabbit anti-LaminB1 1:200 (Abcam Rab16048), mouse anti-Myc 1:200 (Alfagene/Life Technologies #13-2500). The secondary antibodies (1:600 dilution) used were Alexa Fluor 488, Alexa Fluor 555 and Alexa Fluor 647 (Life Technologies). Phalloidin conjugated with Alexa Fluor 488 and Alexa Fluor 647 (Life Technologies) were used to stain actin (1 =200 dilution). DAPI (Sigma-Aldrich) was used to stain the nucleus (1: 10000 dilution).

### Cell imaging

For centrosome reorientation images were acquired in a Zeiss Cell Observer widefield inverted microscope equipped with sCMOS camera Hamamatsu ORCA-flash4,0 V2 for 10ms/frame streaming acquisition, EC Plan-Neofluar 40x/0,75 M27 Oil Objective, LED light source Colibri2 from Zeiss, FS38HE excitation 450-490 nm and emission 500-550 nm and FS43HE excitation 538-562 nm and emission 570-640 nm controlled by with ZEN Blue Edition. For TAN lines quantification the same microscope was used although using a 63x/1.4 Plan-Apochromat DIC M27 Oil objective. Actin retrograde flow analysis was performed using a Zeiss Cell Observer spinning disk confocal inverted microscope equipped with a 37°C chamber, 5% CO_2_ live-cell imaging chamber, Evolve 512 EMCCD camera, confocal scanner Yokogawa CSU-x1, 63x Plan-Apochromat Oil Objective, LED light source Colibri2 from Zeiss, solid state laser 405 nm and 561 nm controlled by ZEN Blue Edition, Confocal images for protein localization at TAN lines were acquired using Zeiss LSM 710 microscope equipped with a 63 /1,4 Plan-Apochromat DIC M27 Oil objective, diode laser 405-30 (405 nm), Argon laser (458, 488 and 514 nm), DPSS 561-10 laser (561 nm) and HeNe633 laser (633 nm).

### Image analysis and figure production

Fiji software was used as an imaging processing software and to quantify actin filaments thickness, Adobe Photoshop and Adobe Illustrator were used to obtain figures.

### Recombinant protein purification

For His-Ctdnep1_Cter production and purification, pDEST42-Ctdnep1_C-ter plasmid was transformed in Rosetta™ 2(DE3)pLysS Competent bacteria (Merck Millipore), Bacteria cultures (500 ml) were grown with the proper antibiotics at 20°C until the optical density (OD) was between 0.6 and 0.8, Protein expression was induced by adding 0.4 mM IPTG and the bacteria cultures were grown for 16-20 hours at 16°C. The bacteria culture was centrifuged 15 minutes at 4000 xg and the pellet was resuspended in 20 ml Ctdnep1 Lysis buffer (50mM Tris-HCl pH 7.5, 200 mM NaCl, lysozyme 0,1 mg/ml, 20 mM imidazole, 1 mM DTT, 0.5 mg/ml Pefabloc^®^ (Roche), 0.5 mM EDTA). The lysate was sonicated on ice for 15 minutes (10 seconds ON, 10 seconds OFF) and centrifuged for 30 minutes, 10000 g at 4°C. The supernatant was incubated with 400 μl of Ni-NTA beads (Life Technologies), previously washed with PBS and lysis buffer, for 1 hour at 4°C and with rotation. Posteriorly, the beads were centrifuged at 800 xg for 4 minutes and washed 2 times with Wash Buffer 1 (50 mM Tris-HCl pH 7.5, 200 mM NaCl, 20 mM Imidazole, 1 mM DTT). The beads were washed 2 times with Wash Buffer 2 (50 mM Tris-HCl pH 7.5, 1 M NaCl, 30 mM Imidazole). The beads were kept in PBS, 1 mM DTT and 40% glycerol at −80°C. To elute His-Ctdnep1_C-ter protein, the beads were incubated for 2 hours at 4°C in Elution Buffer (50 mM Tris-HCL pH 7.5, 200 mM NaCl, 400 mM Imidazole and 1 mM DTT). The eluted fraction was concentrated using a 3 KDa MWCO Amicon^®^ Ultra-15 Centrifugal Filter Unit using Stockage Buffer (50 mM Tris-HCl pH 7.5 and 200 mM NaCl).

For GST, GST-Eps8L2 and GST-Eps8L2 fragments, pDEST15 plasmids were transformed in Rosetta™ 2(DE3)pLysS Competent bacteria (Merck Millipore). Bacteria cultures were grown with the proper antibiotic at 37°C until an OD of 0.6 was obtained. Protein expression was induced adding 0.1 mM IPTG and incubate for 4 hours at 34°C. The bacteria pellet was recovered by centrifuging 15 minutes at 4000 xg. The pellet was lysed with 20 ml of Eps8L2 Lysis Buffer (10 mM Tris.HCLpH8, 150 mM NaCl, 1 mMEDTA, lysozyme 0.1 mg/ml, Pefabloc^®^ (Roche) 0.5 mg/ml) for 30 minutes at 4°C to dissolve the pellet. DTT (1 mM) and Sarkosyl (1.4%, dissolved in Eps8L2 Lysis Buffer without lysozyme nor Pefabloc^®^ (Roche)) were added posteriorly. The lysate was sonicated on ice for 15 minutes (10 seconds ON, 10 seconds OFF) and centrifuged for 45 minutes at 39000 xg and 4°C. The supernatant was collected and incubated with 20 ml of Eps8L2 Lysis Buffer without lysozyme nor Pefabloc^®^ (Roche), and 4% Triton X-100 for 30 minutes at 4°C. The lysate was then incubated with Glutathione Sepharose 4B beads (GE Healthcare Life Sciences) previously washed 3 times with cold PBS. The incubation was for 2 hours at 4°C with rotation. Then, the beads were centrifuged for 4 minutes at 800 xg and washed three times with cold PBS. The beads were kept in PBS, 1 mM DTT and 40% glycerol at −80°C. It is important to mention that the fragment GST-Eps8L2_297-370 was impossible to purify due to its insolubility.

To check the purification process, an aliquot of each step was taken to run a SDS-Page for BlueSafe (NZYTech #MB15201) staining or Western Blot.

### Immunoprecipitation

For *in vitro* Pull Down, 20 μg of GST and GST-Eps8L2 beads (completed with Glutathione Sepharose 4B beads until 30 μl of total beads if needed) were washed with Wash/Binding Buffer (125 mM Tris-HCl, 150 mM NaCl). The beads were incubated with equal amount of eluted His-Ctdnep1_C-ter (final volume of 300 μl in Wash/Binding Buffer) for 3 hours at 4°C with rotation. The beads were posteriorly washed (by centrifuging at 800 xg for 5 minutes) three times with Wash/Binding Buffer. All supernatant was removed using a 1 ml syringe and the beads were resuspended in 30 μl 2X SDS Sample Buffer (Merck Millipore). The samples were incubated at 98°C for 5 minutes before electrophoresis.

For the Eps8L2 fragments Pull Down, 20 ug of GST, GST-Eps8L2 and GST-Eps8L2 fragments proteins bound to glutathione beads (complete with Glutathione Sepharose 4B beads until 30 μl of total beads if needed) were washed with Wash/Binding Buffer. To make GFP-KDEL, Ctdnep1-GFP and Ctdnep1_D67E-GFP lysates, U2OS cells were transfected with the plasmids and lysates were made using U2OS Lysis Buffer (25 mM Tris-HCl, 100 mM NaCl, 1% Triton X-100, 10 % Glycerol). The beads were incubated with 300 μl of lysates for 3 hours at 4°C with rotation. Posteriorly, the beads were washed (by centrifuging at 800 xg for 5 minutes) three times with IP 1500 Buffer (50 mM Tris-HCl pH 7.5, 50 mM NaCl, 0,1% Triton X-100). All supernatant was removed using a 1 ml syringe and the beads were resuspended in 30 μl 2X SDS Sample Buffer (Merck Millipore). The samples were incubated at 98°C for 5 minutes before electrophoresis.

Co-Immunoprecipitation of Ctdnep1 and endogenous Eps8L2 was performed by using SKBR-3 cell lysates (high amount of endogenous Eps8L2 expression). SKBR-3 cells were incubated with SKBR Lysis Buffer (20 mM Hepes pH 7, 10 mM KCl, 0.1% NP40, 6 mM MgCl2, 20% glycerol) on ice for 10 minutes. Lysates were sonicated for 15 seconds and 10 mA and they were incubated with 5U of DNaseI 60 min at 4°C. For pre-clearing, lysates were incubated with 10 μl of Protein A/G magnetic beads (Life Technologies) for 30 minutes at 4°C and rotation. The beads were collected using a magnetic rack. Lysates (1 mg total protein) were incubated with Protein A/G magnetic beads and 2.8 mg of Eps8L2 antibody in IP Buffer (500ul total volume, 20 mM Hepes pH 7, 10 mM KCl, 1.5 mM MgCl2, 0.2% Tween20, 10% glycerol, 1 mM DTT), Incubation was performed overnight at 4°C, The beads were collected using a magnetic rack and washed with Wash Buffer 1 (20 mM Hepes pH 7, 50 mM KCl, 1.5 mM MgCl2, 0.2% Tween20, 10% glycerol, 1 mM DTT), Wash Buffer II (20 mM Hepes pH 7, 100 mM KCl, 1.5 mM MgCl2, 0.2% Tween20, 10% glycerol, 1 mM DTT) and Wash Buffer III (20 mM Hepes pH 7, 150 mM KCl, 1,5 mM MgCl2, 0.2% Tween20, 10% glycerol, 1 mM DTT), Finally, the beads were resuspended in 30 μl 2X SDS Sample Buffer (Merck Millipore) and incubated at 55°c for 5 minutes before electrophoresis.

To immunoprecipitate myc-Eps8L2 for the phosphorylation assay, 500 μl (30 μg/μl total protein) of the different lysates were pre-cleared with 20 μl of agarose beads previously washed two times with PBS and once with U2OS Lysis Buffer. The lysates were collected by centrifuging at 16000 g for 10 minutes and incubated with 50 μl of Myc-Trap^®^_A beads (Chromotek #yta-20, previously washed as before) for 2 hours at 4°C with rotation. The beads were washed three times with IP1500 buffer. The supernatant was removed with a 1 ml syringe and the beads were resuspended in 30 μl 2X SDS Sample Buffer (Merck Millipore). The samples were incubated at 98°C for 5 min and the samples were posteriorly collected by centrifuging for 5 minutes at 16000 xg.

Yeast Two-Hybrid assay was performed by Hybrigenics using Ctdnep1_Cter as bait fragment, Human Placenta_RP5 as prey library and pB27 (N-LexA-bait-C fusion) as vector.

### Phosphorylation assays

To analyse Eps8L2 phosphorylation state, the samples were run in SuperSep™ Phos-tag™ 7.5% (Wako Chemicals) following manufacturer’s instructions. As a control, the same samples were run in parallel in a Mini-PROTEAN^®^ TGX 4-15% Precast protein gels (BioRad). For mass spectrometry assay, U2OS cells were transfected with myc-Eps8L2 and co-transfected with Ctdnep-GFP or Ctdnep1 _D67E-GFP. Lysates were made as indicated in immunoprecipitations section. Immunoprecipitations were performed in three independent expeirments and all the samples were send to Proteomics Core Facility at EMBL (Heidelberg). TMT labelling for the individual samples was performed according to the manufacture’s instructions. Samples preparation before mass spectrometry to identify and quantify Eps8L2 phosphopeptides was performed according to previous work [66].

### Western blot

Protein samples were run in Mini-PROTEAN^®^ TGX 4-15% Precast protein gels (BioRad) and transferred to nitrocellulose membranes. Western blots were probed using the following primary and secondary antibodies (incubated in PBS, 0.1% Tween20 and 5% milk): rabbit anti-Eps8L2 (Sigma-Aldrich #HPA041143, 1:500), mouse anti-Flag (Sigma-Aldrich #F1804, 1:1000), mouse anti-6X His (Abcam Rab18184, 1:1000), chicken anti-GFP (Aves Labs #GFP-1020, 1:2000), anti-mouse HRP (Thermo Scientific #32430, 1:5000), anti-rabbit HRP (Thermo Scientific #31460, 1:5000), anti-chicken HRP (Jackson ImmunoResearch Laboratories # 703-035-155, 1:5000).

### Data analysis

Graphpad Prism 8 software was used to analyse and represent the data. Results are expressed as mean +/− SEM for bars plots. Boxes plots represent 5-95 percentile (outliers are represented as dots/squares) or minimum-maximum values. The box represent the interquartile range (IQR) that includes from 25th percentile (Q1) to 75th percentile (Q3). The line indicate the median value. All quantifications were made with at least three different replicates for experiment. Statistical significance was assessed using unpaired t-tests.

**Table M1.**
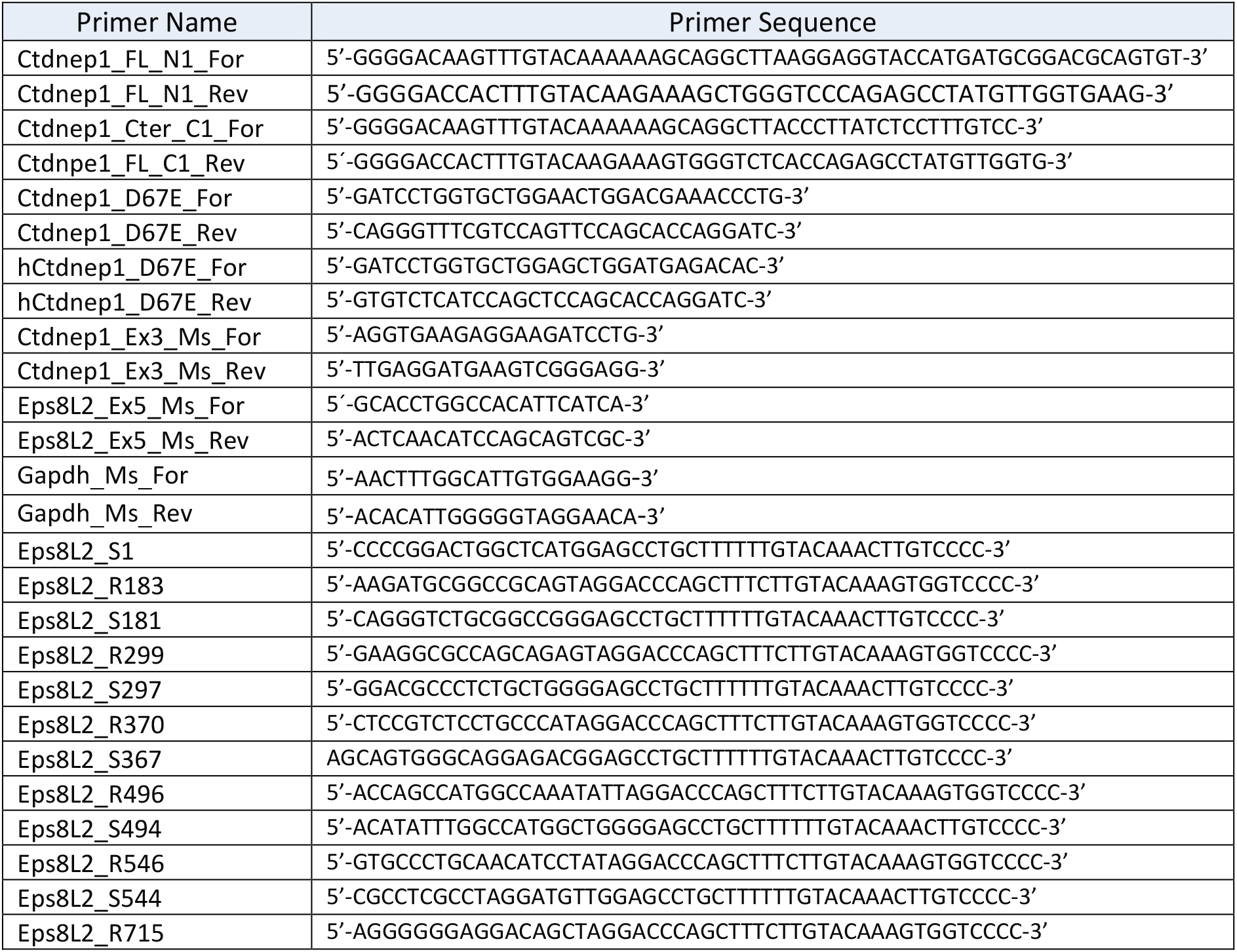
Primers

**Table M2.**
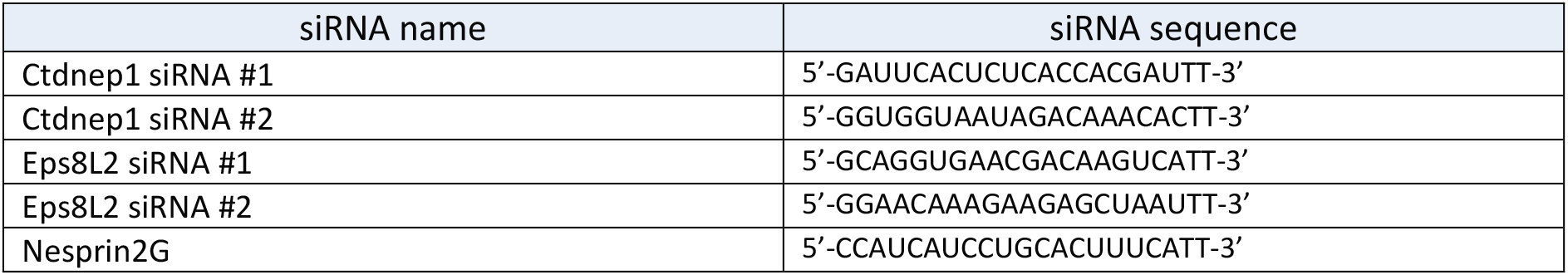
siRNAs

**Table M3.** Plasmids

| Plasmid name | Origin |
| --- | --- |
| pDONR201-Ctdnep1 | This work |
| pDONR201-Ctdnep1_D67E | This work |
| pDONR201-Ctdnep1_C-ter | This work |
| pDEST47-Ctdnep1_WT | This work |
| pDEST47-Ctdnep1_D67E | This work |
| pDEST17-Ctdnep1_C-ter | This work |
| pDONR201-Eps8L2 | This work |
| pDONR201-Eps8L2_1-183 | This work |
| pDONR201-Eps8L2_181-299 | This work |
| pDONR201-Eps8L2_297-370 | This work |
| pDONR201-Eps8L2_367-496 | This work |
| pDONR201-Eps8L2_494-546 | This work |
| pDONR201-Eps8L2_544-715 | This work |
| pDEST15- Eps8L2 | This work |
| pDEST15- Eps8L2_1-183 | This work |
| pDEST15- Eps8L2_181-299 | This work |
| pDEST15- Eps8L2_297-370 | This work |
| pDEST15- Eps8L2_367-496 | This work |
| pDEST15- Eps8L2_494-546 | This work |
| pDEST15- Eps8L2_544-715 | This work |
| pRK5mycGW- Eps8L2 | This work |
| pEZYflag-Eps8L2 | This work |
| pcDNA3.1(+)-C-eGFP- Ctdnep1_WT_siR | This work |
| pcDNA3.1(+)-C-eGFP- Ctdnep1_D67E_siR | This work |
| pDONR201 | Life Technologies |
| pDEST47 | Life Technologies |
| pDEST15 | Fanny Jaulin Lab |
| pRK5mycGW | Fanny Jaulin Lab |
| pEZYflag | Addgene #18700 |
| GFP-NLS | [67] |
| pUBC-GFP-KDel | [65] |
| pDEST17 | Fanny Jaulin Lab |
| pLALI_Lifeact-mcherry | Olivier Pertz |
| pEGFP-C1-GFP-miniNesprin2G | [25] |
| pcDNA3.1-Ctdnep1-2xFlag | [39] |
| pcDNA3.1-Ctdnep1_D67E-2xFlag | This work |
| pGEX-6P-1 | GE Healthcare Life Sciences |

**Table M4.** Antibodies

| Antibody/Probe | Source | Reference | Concentration |
| --- | --- | --- | --- |
| Rabbit anti-β-Catenin | Invitrogen | #712700 | IF→ 1:200 |
| Mouse anti-Pericentrin | BD-Biosciences | #611814 | IF→ 1:200 |
| Mouse anti-Flag | Sigma-Aldrich | #F1804 | IF→ 1:200<br>WB→ 1:1000 |
| Chicken anti-GFP | Aves Lab | #GFP-1020 | IF→ 1:1000<br>WB→ 1:5000 |
| Rabbit anti-LaminB1 | Abcam | #ab16048 | IF→ 1:200 |
| Mouse anti-Myc | Life Technologies | #13-2500 | IF→ 1:200<br>WB→ 1:1000 |
| Mouse anti-6x His | Abcam | #ab18184 | WB→ 1:1000 |
| Goat anti-Chicken IgY (H+L) Secondary Antibody, Alexa Fluor® 488 conjugate | ThermoFisher Scientific | #A-11039 | IF→ 1:600 |
| Donkey Anti-Mouse IgG (H+L) Highly Cross-Adsorbed Secondary Antibody, Alexa Fluor® 488 | ThermoFisher Scientific | #A21202 | IF→ 1:600 |
| Donkey Anti-Rabbit IgG (H+L) Highly Cross-Adsorbed Secondary Antibody, Alexa Fluor® 488 | ThermoFisher Scientific | #A21206 | IF→ 1:600 |
| Goat anti-Mouse IgG (H+L) Secondary Antibody, Alexa Fluor® 555 conjugate | ThermoFisher Scientific | #A21424 | IF→ 1:600 |
| Goat Anti-Rabbit IgG (H+L) Highly Cross-Adsorbed Secondary Antibody, Alexa Fluor® 555 | ThermoFisher Scientific | #A21429 | IF→ 1:600 |
| Goat anti-Rabbit IgG (H+L) Secondary Antibody, Alexa Fluor® 647 conjugate | ThermoFisher Scientific | #A-21245 | IF→ 1:600 |
| Goat anti-Mouse IgG (H+L) Cross-Adsorbed Secondary Antibody, Alexa Fluor® 647 | ThermoFisher Scientific | #A-21236 | IF→ 1:600 |
| Alexa Fluor® 488 Phalloidin | Life Technologies | #A12379 | IF→ 1:200 |
| Alexa Fluor® 647 Phalloidin | Life Technologies | #A22287 | IF→ 1:100 |
| DAPI | Sigma-Aldrich | #32670-5MG-F | IF→ 1:10000 |

## Acknowledgments

We thank Fanny Jaulin, Shirim Bahmanyar, Jan Lammerding, Gregg Gundersen and Olivier Pertz laboratories for reagents. We acknowledge all members from Edgar Gomes, Claudio Franco, Vanessa Morais and Sergio Almeida laboratory for discussions and support, the Bioimaging facility and all iMM community as well as the Proteomics Core Facility at EMBL, This work was supported by the European Research Council, EMBO installation, Association pour la Recherche sur le Cancer, France and La Ligue Nacionale contre le Cancer (France), LISBOA-01-0145-FEDER-007391 project co-funded by FEDER, through POR Lisboa 2020 - Programa Operacional Regional de Lisboa, PORTUGAL 2020, and Fundação para a Ciência e a Tecnologia.

## Author Contributions

F.J.C-C., D.O., S.C.S., B.C. and E.R.G. conceived and designed the experiments. F.J.C-C., D.O., S.C.S., Y.J., J.D. and S.C.M. performed experiments. F.J.C-C., D.O., and S.C.S. analysed the data, F.J.C-C. and E.R.G. wrote the manuscript. All the authors participated in the critical review and revision of the manuscript.

