## Supplemental Figures for "Ctdnep1 and Eps8L2 regulate dorsal actin cables for nuclear positioning during cell migration"

A

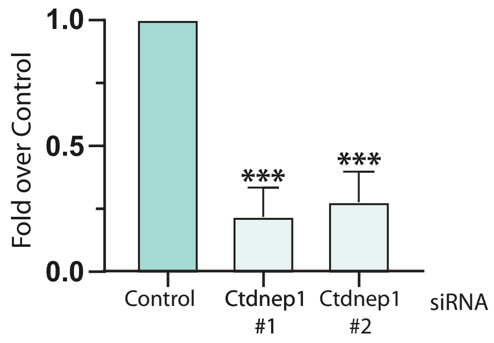


B

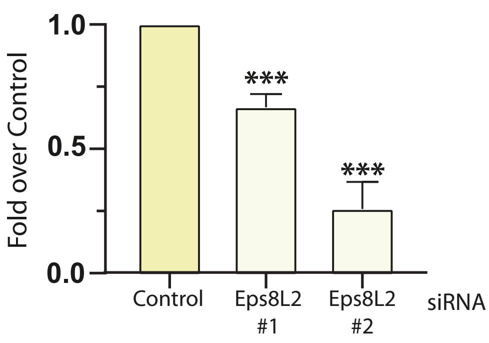


C

**Yeast-2-Hybrid screen main interactions:**

| Clone Name | Gene Name | %Id5p | %Id3p | PBS |
|---|---|---|---|---|
| pB27_A-38 | ADRBK1 | 99.8 | 99.8 | D |
| pB27_A-28 | C18orf24 | 98.6 | 98.6 | D |
| pB27_A-39 | CDCA3 | 100 | 99.8 | A |
| pB27_A-107 | COPE | 99.6 | 99.6 | D |
| pB27_A-77 | CSH1 | 99.8 | 99.4 | D |
| pB27_A-62 | DDX20 | 98.6 | | D |
| pB27_A-42 | EFEMP1 | 100 | 100 | D |
| pB27_A-95 | EPS8L2 | 99.7 | 99.7 | B |
| pB27_A-89 | MGC12987 | 100 | 98.9 | D |
| pB27_A-24 | MKNK2 | 100 | 99.9 | D |
| pB27_A-2 | MKRN1 | 93.7 | 92.3 | D |
| pB27_A-45 | NAIP | 97.5 | 98 | D |
| pB27_A-71 | OSBPL2 | 99.3 | 99.6 | C |
| pB27_A-21 | PARN | 100 | 99.1 | D |
| pB27_A-86 | PDIK1L | 99.4 | 99.4 | D |
| pB27_A-11 | THAP4 | 99.1 | 99.1 | D |
| pB27_A-15 | ZDHHC17 | 100 | 100 | D |

A Very high confidence in the interaction

B High confidence in the interaction

C Good confidence in the interaction

D Moderate confidence in the interaction

**Bait** → Ctdnep1_C-ter

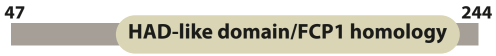


**Prey Library** → Human Placenta_RP5

Calero-Cuenca et al. Figure S1

**Figure S1**. (**A**) Ctdnep1 mRNA quantification in 3T3 fibroblasts transfected with Control, Ctdnep1 #1 and Ctdnep1 #2 siRNAs measured by RT-qPCR. (**B**) Eps8L2 mRNA quantification in 3T3 fibroblasts transfected with Control, Eps8L2 #1 and Eps8L2 #2 siRNAs measured by RT-qPCR. (**C**) Yeast Two-Hybrid screening performed with Ctdnep1_C-ter as the bait and human placenta library as prey. The table shows the main interactions observed in the screening. Each interaction present a Predicted Biological Score (PBS) that is computed to assess the interaction reliability. This score represents the probability of an interaction to be non-specific: it is an e-value, primarily based on the comparison between the number of independent prey fragments found for an interaction and the chance of finding them at random (background noise). %Id5p and %Id3p indicates the % identity of the prey fragment sequences with the gene reference sequence.

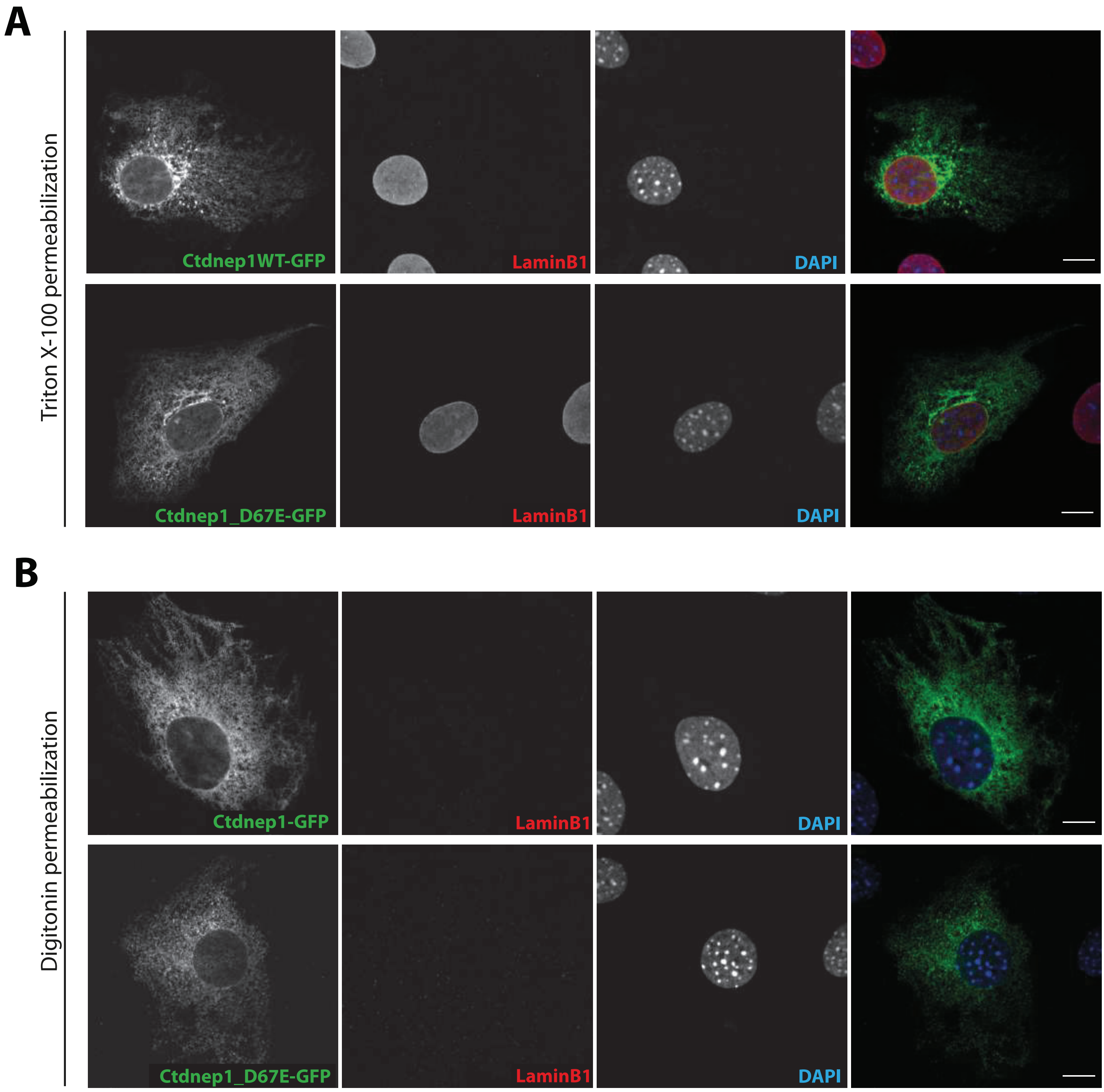


Calero-Cuenca et al. Figure S2

**Figure S2.** Representative image of wound-edge fibroblasts stimulated with LPA and microinjected with Ctdnep1-GFP or Ctdnep1_D67E-GFP fixed and permeabilized with Triton X-100 (**A**) or digitonin (**B**) and stained for GFP (green), laminB (red) and DAPI (blue, nucleus). Triton treatment permeabilizes the plasma membrane and nuclear envelope whereas digitonin only permeabilizes the plasma membrane and not the nuclear envelope. Scale bar: 10 µm. Data are represented as mean +/- SEM.

A

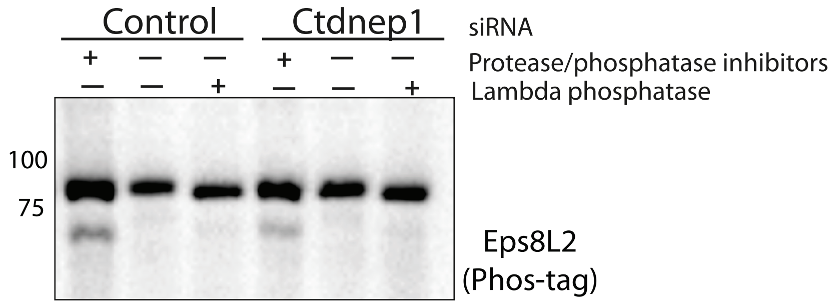


B

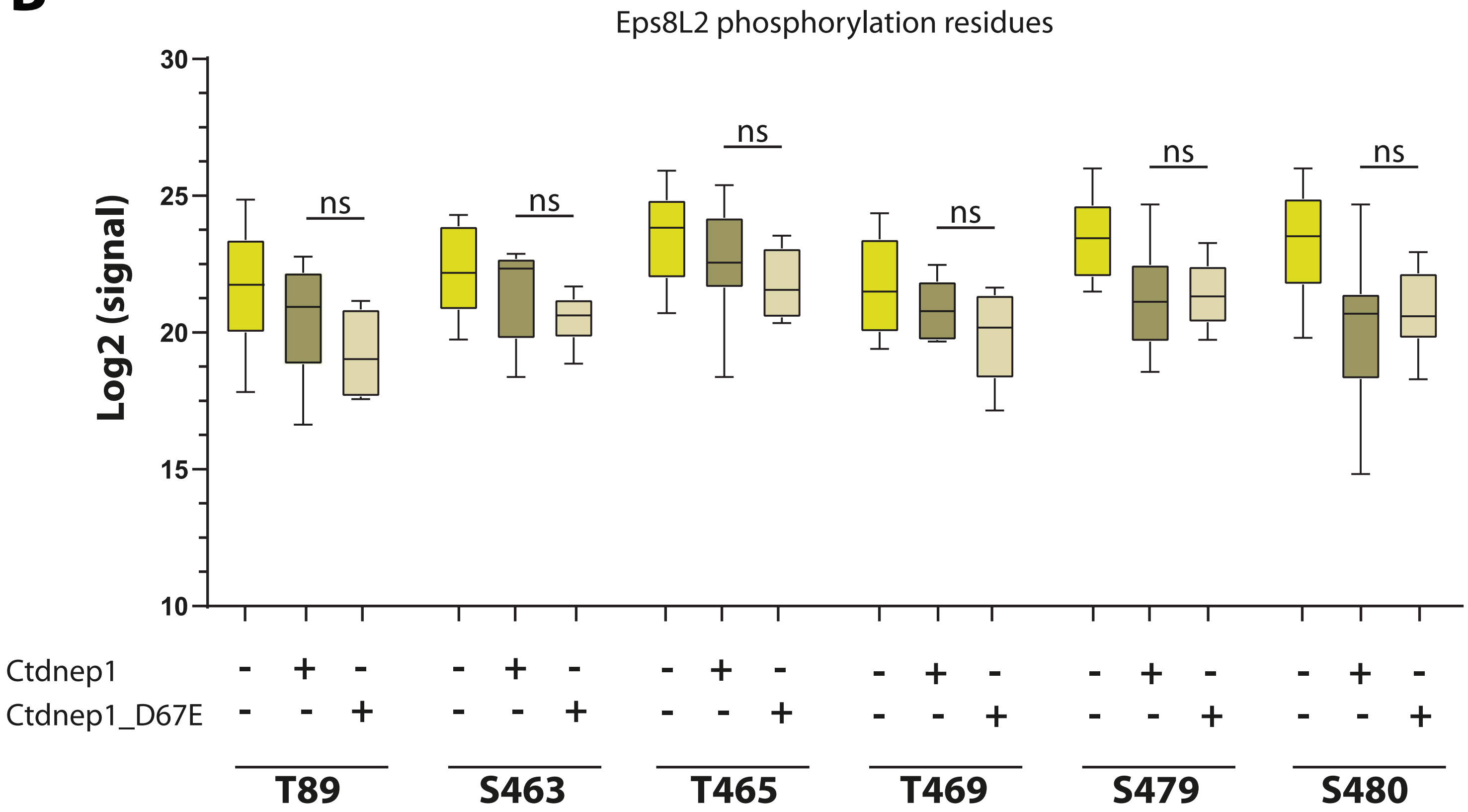


Calero-Cuenca et al. Figure S3

**Figure S3**. (**A**) Phos-tag gel for endogenous Eps8L2 in SKBR-3 cells (that express high Eps8L2) treated with Control and Ctdnep1 siRNAs. Incubation with Lambda phosphatase was used as a control to remove all phosphorylation in the sample. (**B**) Plot representing the $Log_2$ of the signal obtained in the mass spectrometry analysis for each phosphorylated residues detected in Eps8L2 with or without co-transfection with Ctdnep1 and Ctdnep1_D67E.

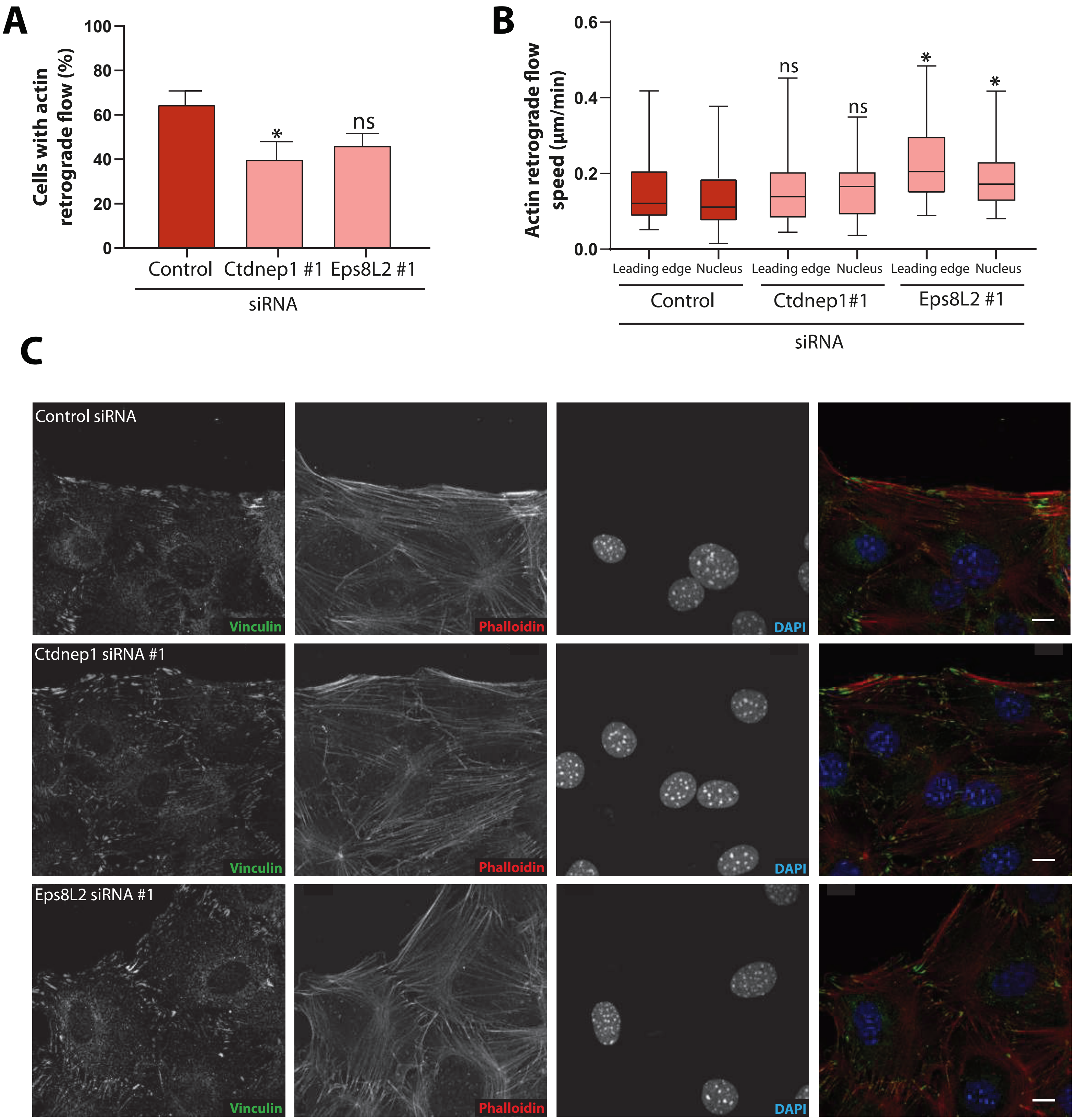


Calero-Cuenca et al. Figure S4

**Figure S4**. (**A**) Quantification of the percentage of wound-edge fibroblasts with actin retrograde flow after LPA stimulation in Control, Ctdnep1 and Eps8L2 siRNAs. A stable cell line of 3T3 fibroblasts expressing lifeact-mcherry were used to measure actin retrograde flow. Data are represented as mean +/- SEM. (**B**) Quantification of actin retrograde flow speed for actin filaments located near the leading edge (Leading edge) or on the dorsal side of the nucleus (Nucleus) in the conditions showed in A. Data are represented as minimum-maximum values. (**C**) Representative images of of wound-edge fibroblasts stimulated with LPA and treated with Control, Ctdnep1 and Eps8L2 siRNAs. Cells were stained for Vinculin (focal adhesions, green), phalloidin (Actin, red) and DAPI (Nucleus, blue). Scale bar: 10 µm. Data are represented as minimum-maximum values.

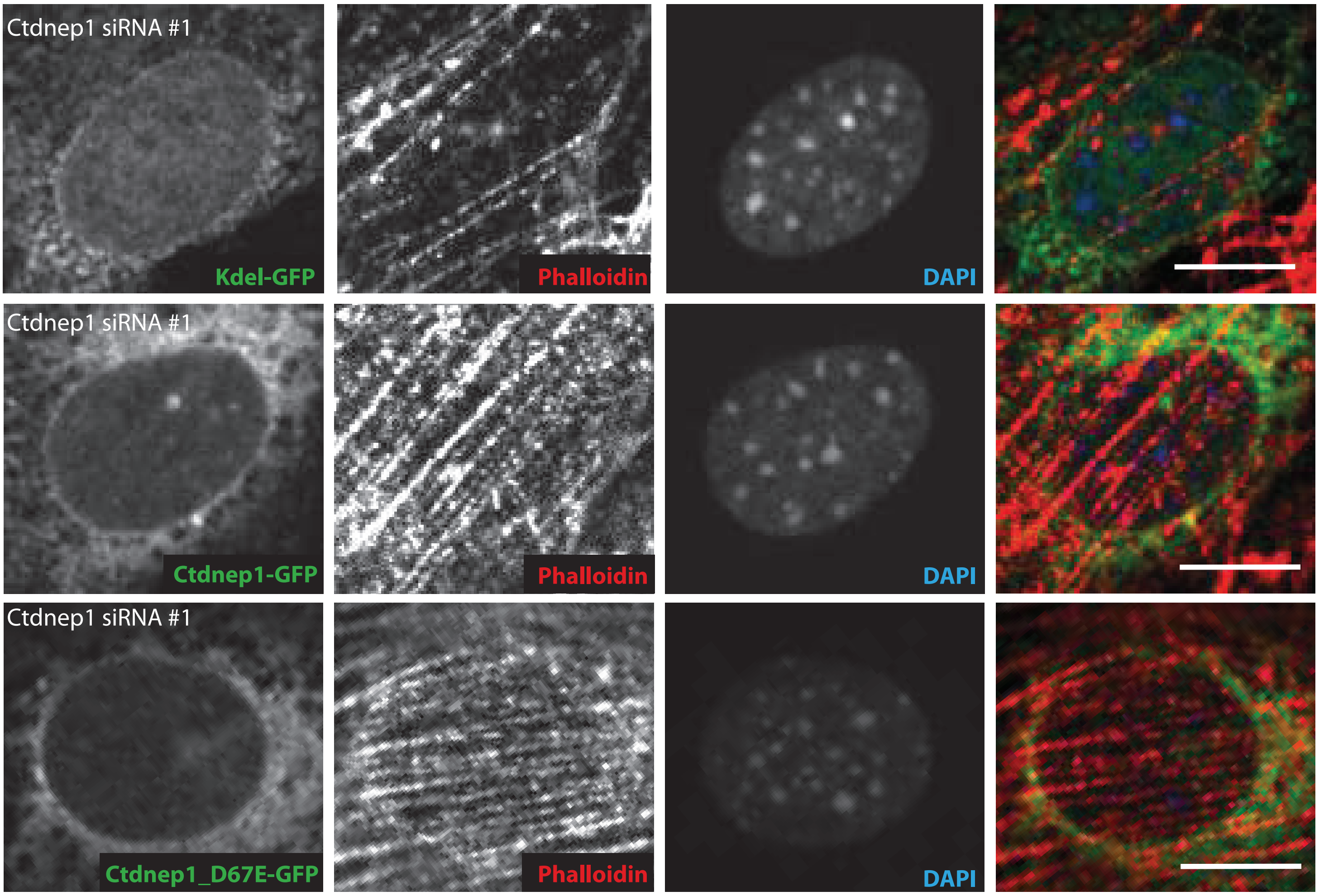


Calero-Cuenca et al. Figure S5

**Figure S5**. (**A**) Representative images of the nucleus of wound-edge fibroblasts stimulated with LPA, treated with Ctdnep1 siRNA and microinjected with KDEL-GFP, Ctdnep1-GFP and Ctdnep1D67E-GFP. Cells were stained for GFP (green), phalloidin (Actin, red) and DAPI (Nucleus, blue). Scale bar: 10 µm.
